# Niche and neutral processes both shape community structure in parallelized, aerobic, single carbon-source enrichments

**DOI:** 10.1101/154781

**Authors:** Theodore M. Flynn, Jason C. Koval, Stephanie M. Greenwald, Sarah M. Owens, Kenneth M. Kemner, Dionysios A. Antonopoulos

## Abstract

Here we seek to test the extent to which laboratory enrichments mimic natural community processes and the degree to which the initial structure of a community determines its response to a press disturbance via the addition of environmentally-relevant carbon compounds. By utilizing aerobic substrate arrays to examine the effect of carbon amendment on microbial communities taken from six distinct environments (soil from a temperate prairie and forest, tropical forest soil, subalpine forest soil, and surface water and soil from a palustrine emergent wetland), we examined how carbon amendment and inoculum source shape the composition of the community in each enrichment. Dilute subsamples from each environment were used to inoculate 96-well microtiter plates containing triplicate wells amended with one of 31 carbon sources from 6 different classes of organic compound (phenols, polymers, carbohydrates, carboxylic acids, amines, amino acids). After incubating each well aerobically in the dark for 72 hours, we analyzed the composition of the microbial communities on the substrate arrays as well as the initial inocula by sequencing 16S rRNA gene amplicons using the Illumina MiSeq platform. Comparisons of alpha and beta diversity in these systems showed that, while the composition of the communities that grow to inhabit the wells in each substrate array diverges sharply from that of the original community in the inoculum, these enrichment communities are still is strongly affected by the inoculum source. We found most enrichments were dominated by one or several OTUs most closely related to aerobes or facultative anaerobes from the *Proteobacteria* (e.g. *Pseudomonas*, *Burkholderia*, and *Ralstonia*) or *Bacteroidetes* (e.g. *Chryseobacterium*). Comparisons within each substrate array based on the class of carbon source further show that the communities inhabiting wells amended with a carbohydrate differ significantly from those enriched with a phenolic compound. Niche selection therefore seems to play a strong role in shaping the communities in the substrate arrays, although some stochasticity is seen whereby several replicate wells within a single substrate array display strongly divergent community compositions. Overall, the use of highly parallel substrate arrays offers a promising path forward to study the response of microbial communities to a changing environment.

## 1 Introduction

From the soil under our feet to the deepest sedimentary basins, microbial life inhabits nearly every environment on Earth (Whitman et al., 1998). The abundance and activity of the individual populations that comprise these communities change dynamically in response to changes in their local environment (Nemergut et al., 2013). The composition of microbial communities has been linked to specific parameters like water chemistry in environments such as lakes (Youngblut et al., 2014), streams (Zeglin, 2015), wetlands (Baldwin et al., 2006; Peralta et al., 2010; Dalcin Martins et al., 2017), and aquifers (Flynn et al., 2013; Hug et al., 2015; Kirk et al., 2015). In seawater, for example, the abundance of individual populations of bacteria have been shown to oscillate in sync with changes in light, temperature, and salinity (Eren et al., 2013; Ottesen et al., 2014). Soil, however, possesses such extreme physical, chemical, and biological heterogeneity that understanding the environmental forces that shape the structure and function of microbial communities there remains an outstanding challenge (Tiedje et al., 1999; Roesch et al., 2007; O’Brien et al., 2016; Bailey et al., 2017).

As the number of taxonomic groups within a particular soil often exceeds several thousand distinct clades, there is considerable interest in using simplified microbial communities to test hypotheses related to soil ecology. By “minimizing” a native microbial community from soil or elsewhere through enrichment in the laboratory, noise from the myriad co-existing metabolic networks and structural heterogeneities present in the parent environment can be pared down to focus on a particular process of interest, allowing the power of modern omics technology to be brought to bear on specific questions in microbial ecology (Prosser, 2015). This microcosm approach is frequently used to examine a subset of a native community such as sulfate reducers (Raskin et al., 1996; Kirk et al., 2013; Kwon et al., 2014), denitrifiers (Laverman et al., 2010; Kraft et al., 2014), or organisms capable of degrading a specific compound of interest (Brennan et al., 2006; Sutton et al., 2013; Luo et al., 2014; Onesios-Barry et al., 2014). Conducting such experiments as high-throughput, parallel replicates across a broad variety of environments has the potential to provide greater insight into the selective and stochastic ecological forces that guide community assembly.

Physiological profiling using microtiter plates has been frequently used as a method of examining the functional diversity of mixed microbial communities by monitoring the production of NADH using a redox-active dye (Bochner, 1989; Bochner et al., 2001). This approach allows the utilization of an array of carbon compounds by microbial communities to be monitored in parallel using 96-well plates. These substrate arrays are frequently used to test the metabolic capabilities of the microbiome inhabiting soil, water, and other environments (Bartscht et al., 1999; King, 2003; Weber and Legge, 2009; Gryta et al., 2014; Zhang et al., 2014). Given that the composition of the communities that grow from the inocula in these arrays is rarely, if ever, characterized directly, the extent to which the “active” populations utilizing a particular substrate are representative of the community at large remains unclear (Konopka et al., 1998; Haack et al., 2004). Substrate arrays also offer an opportunity to study in parallel the response of a single inoculum to the addition of a wide array of nutrients.

In this study, we employed these substrate arrays as mini-bioreactors to monitor the response of microbial communities from six distinct environments to enrichment on 31 individual carbon compounds. We used as inocula soil from a wetland, a grassland, and three types of forest (temperate, subalpine, and tropical) as well as surface water from the wetland. We sought to test A) the extent to which communities enriched in these substrate arrays are representative of the initial community as a whole, B) the degree to which the initial structure of a community determines its response to a press disturbance (increase in nutrient levels) for different classes of environmentally-relevant (Hitzl et al., 1997) carbon compounds and C) whether the composition of the communities that grow to inhabit the substrate arrays are more influenced by the composition of the inoculum or the type of carbon source upon which they are enriched.

## 2 Materials and Methods

### 2.1 Experimental Setup

Substrate array experiments were conducted by inoculating 96-well plates with soil suspensions or natural waters from six distinct environments. For these substrate arrays we chose the EcoPlate array (Biolog, Hayward, CA). Each well of this substrate array contains a proprietary minimal media (see Bochner et al. (2001)) as well as one of 31 carbon sources representing five distinct categories: amines, amino acids, carbohydrates, polymers, and phenolic compounds (Table 1). While an exact formulation of the media is not publicly available, the manufacturer’s documentation states that medium components are present at concentration of 2-20 mM for the carbon source, 1-5 mM for N, 0.1-1 mM for P, 0.1-1 mM for S, and < 2 μM of a vitamin solution.

**Table 1.**
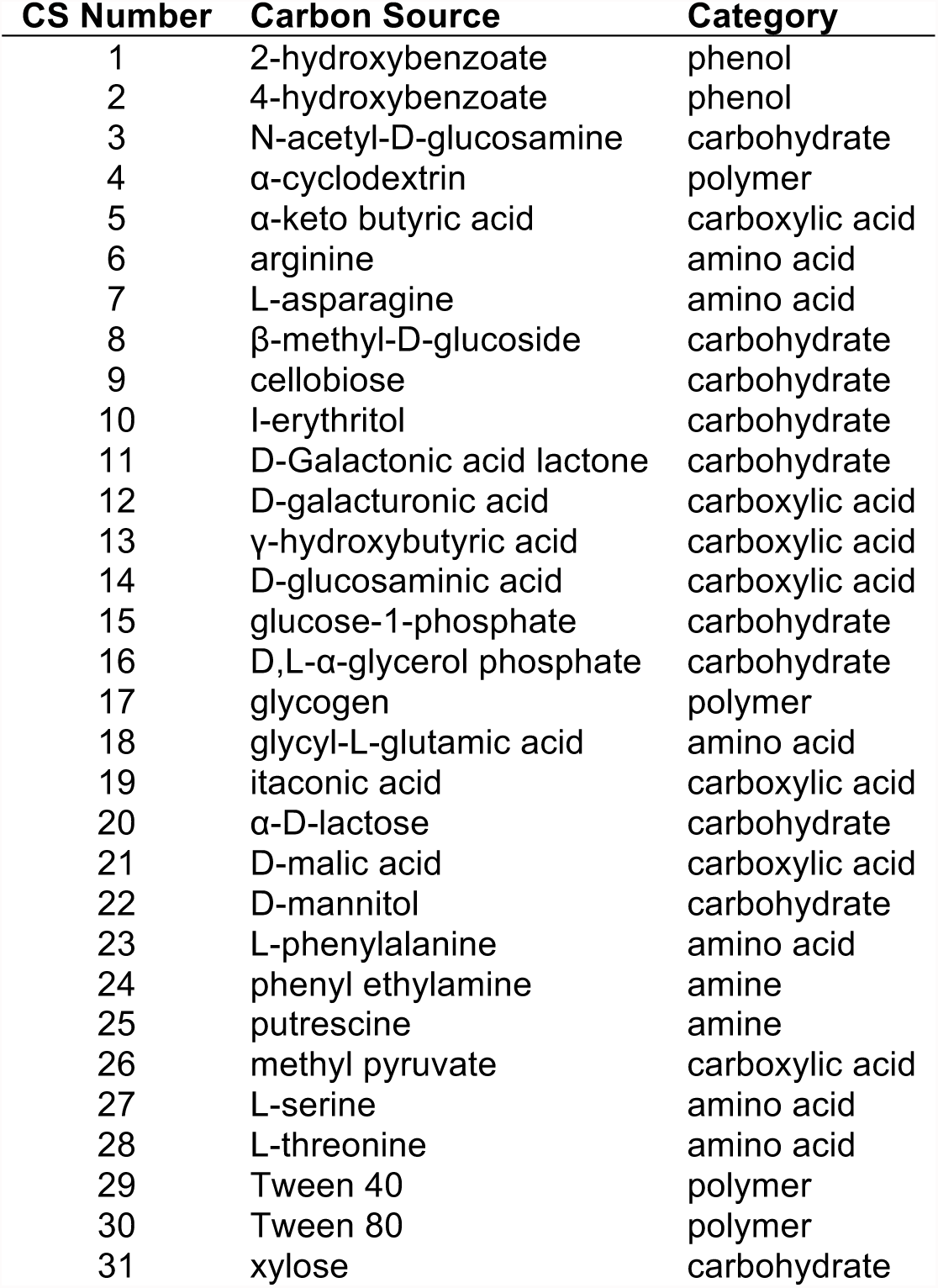
List of 31 carbon substrates present in substrate arrays (1 substrate per well, 3 replicate wells per substrate).

Soil from a temperate climate was collected from a restored prairie grassland (FLP) and an adjacent forest (FLF) on site at the Fermi National Accelerator Laboratory (Fermilab) in Batavia, Illinois, USA. Surface water (AWW) and soil (AWS) was obtained from a palustrine emergent wetland populated by *Typha* and *Phragmites* spp. on site at Argonne National Laboratory in Lemont, Illinois, USA. Tropical forest soil (CRP14) was obtained from a Caribbean lowland rainforest within the EARTH University Forest Reserve in Costa Rica (Alvarez-Clare et al., 2013). Subalpine forest soil (SodaSpr) was taken from a pine forest near Soda Springs, California, USA, at an elevation of 2,063 meters. All soil samples were taken by removing the top 5 cm of soil after clearing off any surface litter. Aquatic samples were taken at the water’s surface using a sterile container. Samples were processed within 72 hours of collection to minimize storage effects.

A conceptual diagram of our experimental design is shown in Figure 1. Solid suspensions used to inoculate the bioreactors were prepared by adding 1 gram of soil to 5 mL of sterile, DNA-free ultrapure water (HyClone HyPure molecular grade, Thermo Scientific). Suspensions were then homogenized using an ultrasonic dismembrator at low intensity for 30 seconds, breaking up any aggregates and removing bacteria from organic and mineral particles. The slurry was then diluted 1:100 (v/v) by adding additional sterile water and dispensed into a substrate array (150 μL per well). When using an aqueous medium as an inoculum in the case of the wetland surface water, the inoculum was amended directly to the array without dilution. After 72 hours of incubation, 100 μL aliquots were taken from each well and used for microbial community analysis by sequencing 16S rRNA gene amplicons.

**Figure 1.**
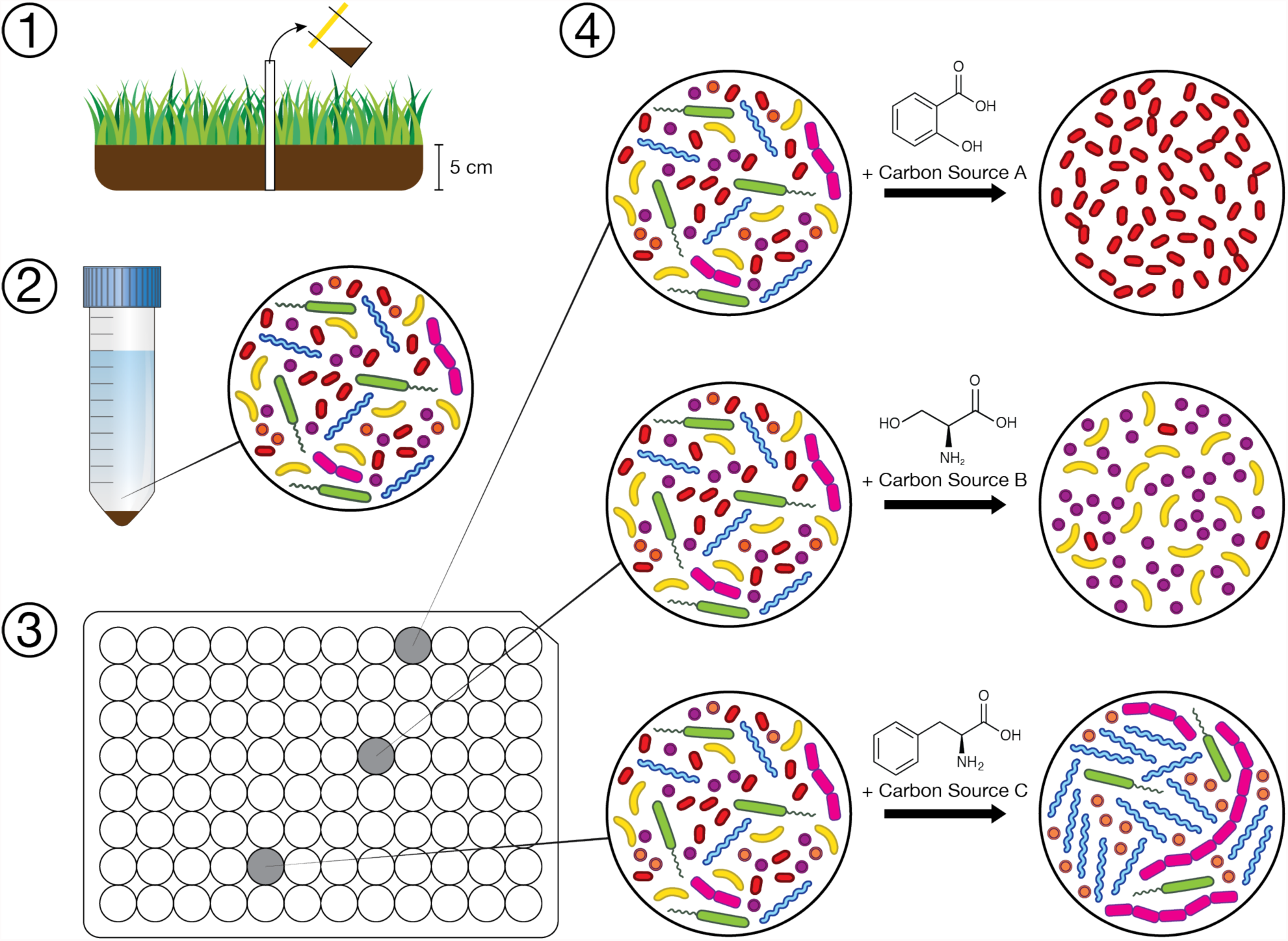
Conceptual diagram of our experimental workflow. 1) Collection of inocula for substrate arrays 2) Dilute suspension of soil/water microbes prepared within 72 hours 3) Substrate arrays where each well is amended with one of 31 individual carbon sources (in triplicate) is inoculated with suspension (1:100 m/v dilution) and incubated for 72 hours 4) Changes in composition of microbial communities in the substrate array is assayed by sequencing 16S rRNA gene amplicons on the Illumina MiSeq platform.

### 2.2 Molecular Microbiology and Bioinformatics

DNA was extracted from both environmental inocula and aliquots from the substrate arrays using MoBio PowerSoil kits following the manufacturer’s instructions. The concentration of DNA extracted was quantified using Quant-iT PicoGreen dsDNA assay (Invitrogen). 16S rRNA genes from each of these samples were amplified by the polymerase chain reaction (PCR) using a primer set (515F-806R) targeting the V4 region of this gene in both bacteria and archaea (Bates et al., 2011; Caporaso et al., 2011). These primers were barcoded to allow for sample multiplexing on the Illumina MiSeq (Caporaso et al., 2012). PCR reactions were carried out using 5 PRIME MasterMix (Gaithersburg, MD). PCR conditions used an initial denaturation step of 95 °C for 3 minutes followed by 35 cycles of 95 °C for 30 seconds, 55 °C for 45 seconds, then 72 °C for 1.5 minutes and finalized by a single extension step 72 °C for 10 minutes. Pooled product for each sample was then quantified using the PicoGreen assay. Primer dimers were removed using the UltraClean PCR Clean-Up Kit (MoBio Laboratories, Inc.) and the amount of DNA in each sample was normalized to a final concentration of 2 ng μL^−1^. Paired-end amplicons (151×12×151 bp) were sequenced on an Illumina MiSeq at the Environmental Sample Preparation and Sequencing Facility at Argonne National Laboratory following procedures described in Caporaso et al. (2012).

A total of 19.0 million paired-end sequences were generated (32,552 ± 20,696 per amplicon library) and then merged followed by downstream processing using QIIME (Caporaso et al., 2010). Briefly, singletons were removed, de novo OTU picking with uclust was used to cluster sequences at 97% similarity, and Greengenes (4feb2011) was used to assign taxonomies. Beta diversity comparisons were conducted in QIIME, Primer-7 (Clarke and Warwick, 2001), or in R using the packages phyloseq (McMurdie and Holmes, 2013) and edgeR (Robinson et al., 2010). Raw sequence data will be freely available to download through MG-RAST (Meyer et al., 2008) upon final publication.

## 3 Results

Parallel substrate arrays containing one of 31 carbon compounds (Table 1) were inoculated with diluted material from one of six different environments: prairie grassland soil (FLP), temperate forest soil (FLF), freshwater wetland surface water (AWW) and soil (AWS), tropical forest soil (CRP14), or subalpine forest soil (SodaSpr). These substrate arrays were incubated at 30 °C for 72 hours, after which time DNA was extracted from the cells present in each of the wells and 16S rRNA genes were sequenced using an Illumina MiSeq.

We observed that, following this incubation, the structures of the microbial communities in the wells of each substrate array deviated sharply from the initial composition of the inoculum. As shown in Figure 2, the average alpha diversity as measured by the Shannon index decreased sharply in the enrichments of all six substrate arrays, falling substantially below the average diversity of the inocula. Nonmetric multidimensional scaling (NMDS) analysis found large differences in the composition of these communities compared to the inoculum based on the weighted UniFrac distance (Figure S1). While the microbial communities in each well of the substrate array diverged from the parent material, they differed substantially from one another as well (Figure 3). The degree of differences between substrate arrays largely recapitulate the observed differences between inocula (Figure S2), with one notable exception. The most divergent communities are from substrate arrays inoculated with soil from the SodaSpr and CRP14 sites. These enrichments differ substantially both from the arrays inoculated with material from temperate environments (AWW, AWS, FLF, and FLP) and from each other. Within those inoculated with materials from Illinois, the FLF and FLP substrate array communities overlap almost entirely. AWS and AWW communities are more separated from each other, but still cluster together with the enrichments inoculated with other Illinois materials (Figure 3).

**Figure 2.**
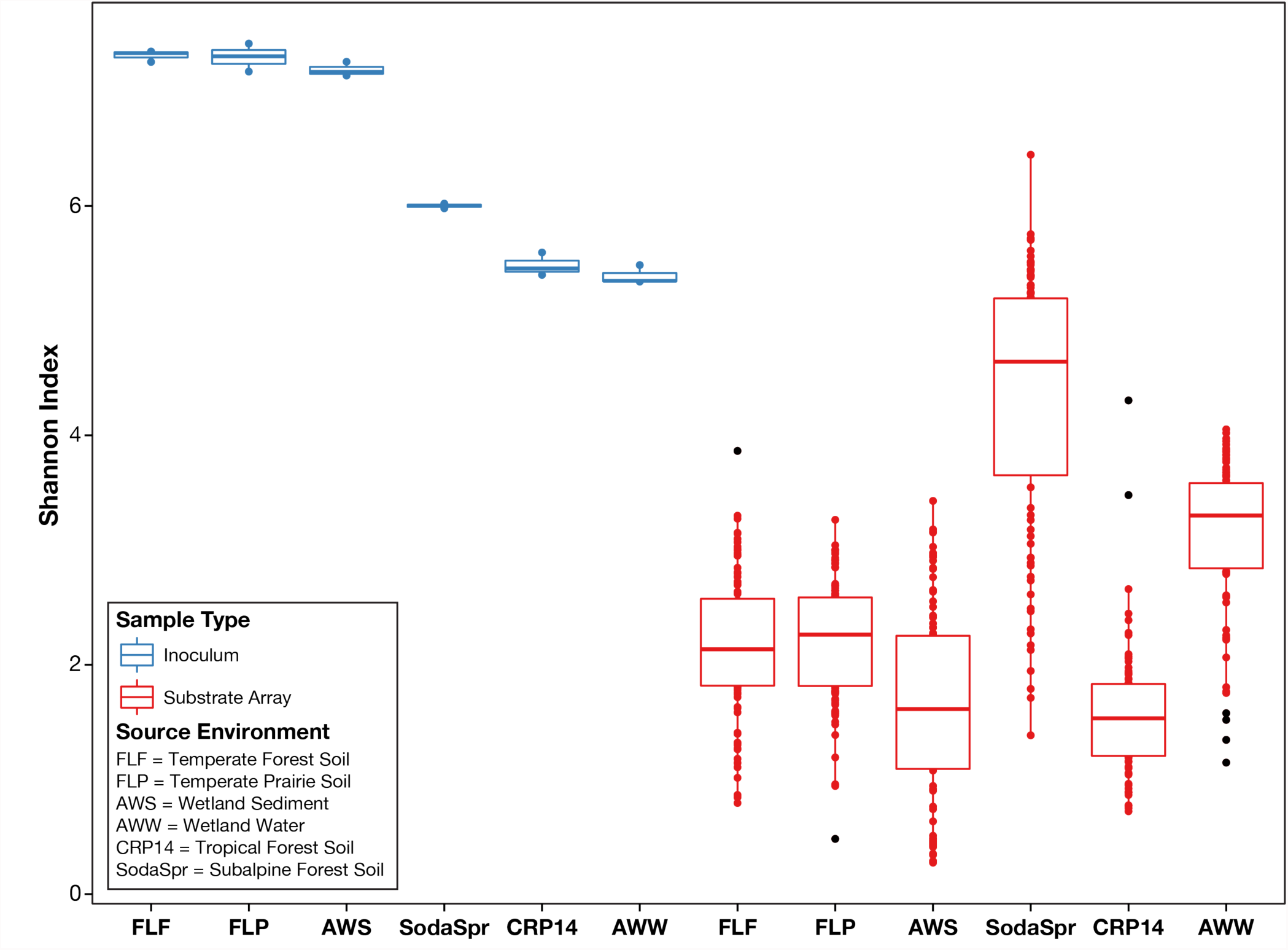
Alpha diversity of environmental inocula and EcoPlate enrichments as measured by the Shannon index of diversity. On average, the total diversity of environmental communities decreases substantially following incubation in the substrate array, regardless of carbon source.

**Figure 3.**
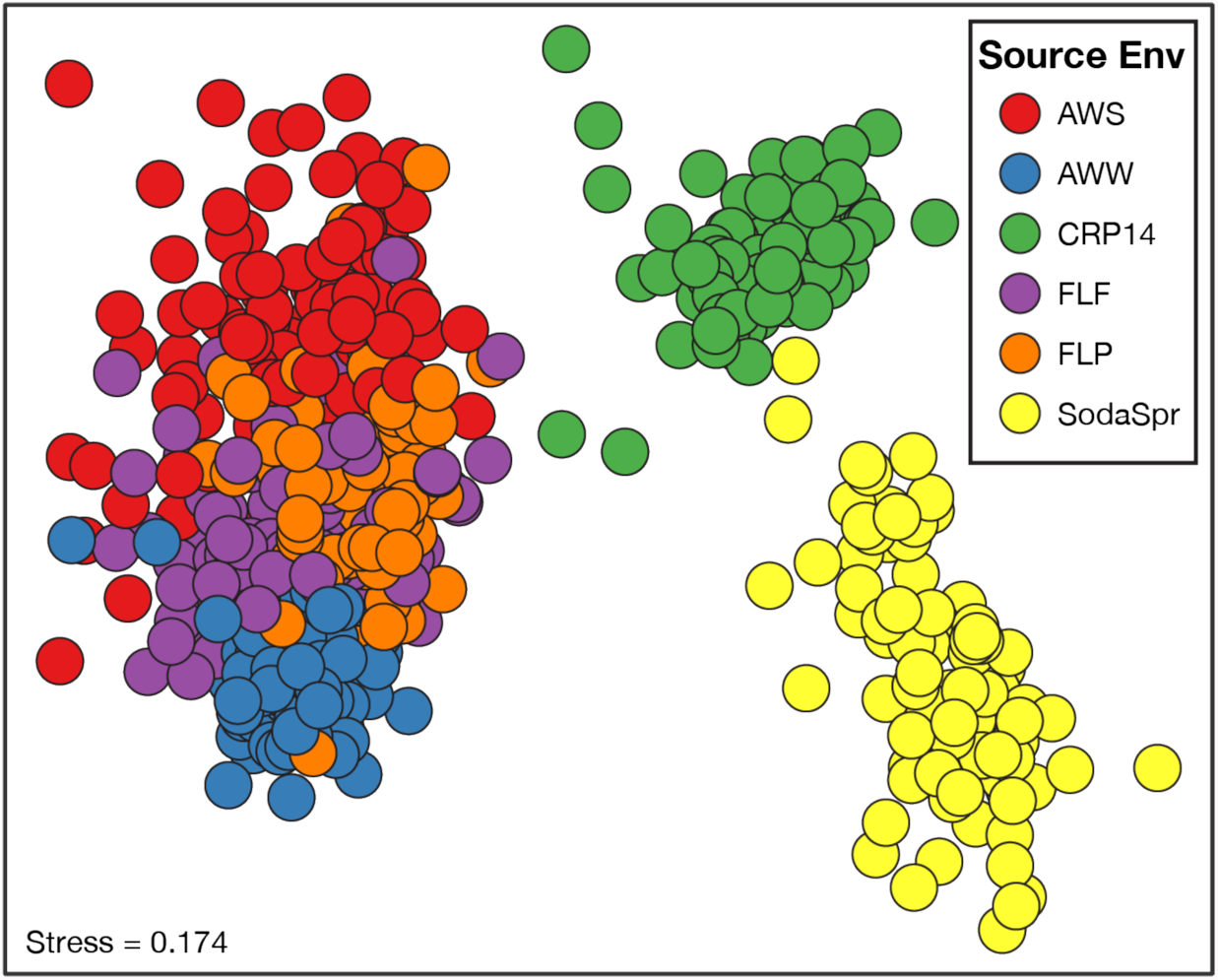
Nonmetric multidimensional scaling (NMDS) analysis based on the weighted UniFrac distance metric showing the difference in microbial community structure between the substrate array enrichments.

Based on analysis of similarity (ANOSIM) shown in Figure 4, the clustering by environment observed in the NMDS ordination shown in Figure 3 is statistically significant for pairwise comparisons between nearly all 6 substrate arrays. The CRP14 communities differ the most from the other substrate arrays, with R-values (R_ANOSIM_) ranging from 0.685-0.876 (*p* < 0.0001%). An R_ANOSIM_ < 0.75 indicates two groups of microbial communities are almost entirely distinct, while a value between 0.25 and 0.75 indicates some degree of overlap (Ramette 2007). Values of R_ANOSIM_ < 0.25 indicate the two groups of communities being compared are largely indistinguishable. Based on this metric, CRP14 enrichments are most similar to (although still mostly distinct from) the SodaSpr enrichments (R_ANOSIM_ = 0.685). SodaSpr enrichments fall on a gradient of distinction between the tropical and temperate enrichments (R_ANOSIM_ = 0.377 -0.685), while temperate soil enrichments show almost complete overlap with one another (R_ANOSIM_ = 0.136-0.288). The two wetland substrate arrays (AWW and AWS) are distinct from one another but share some overlap (R_ANOSIM_ = 0.408).

**Figure 4.**
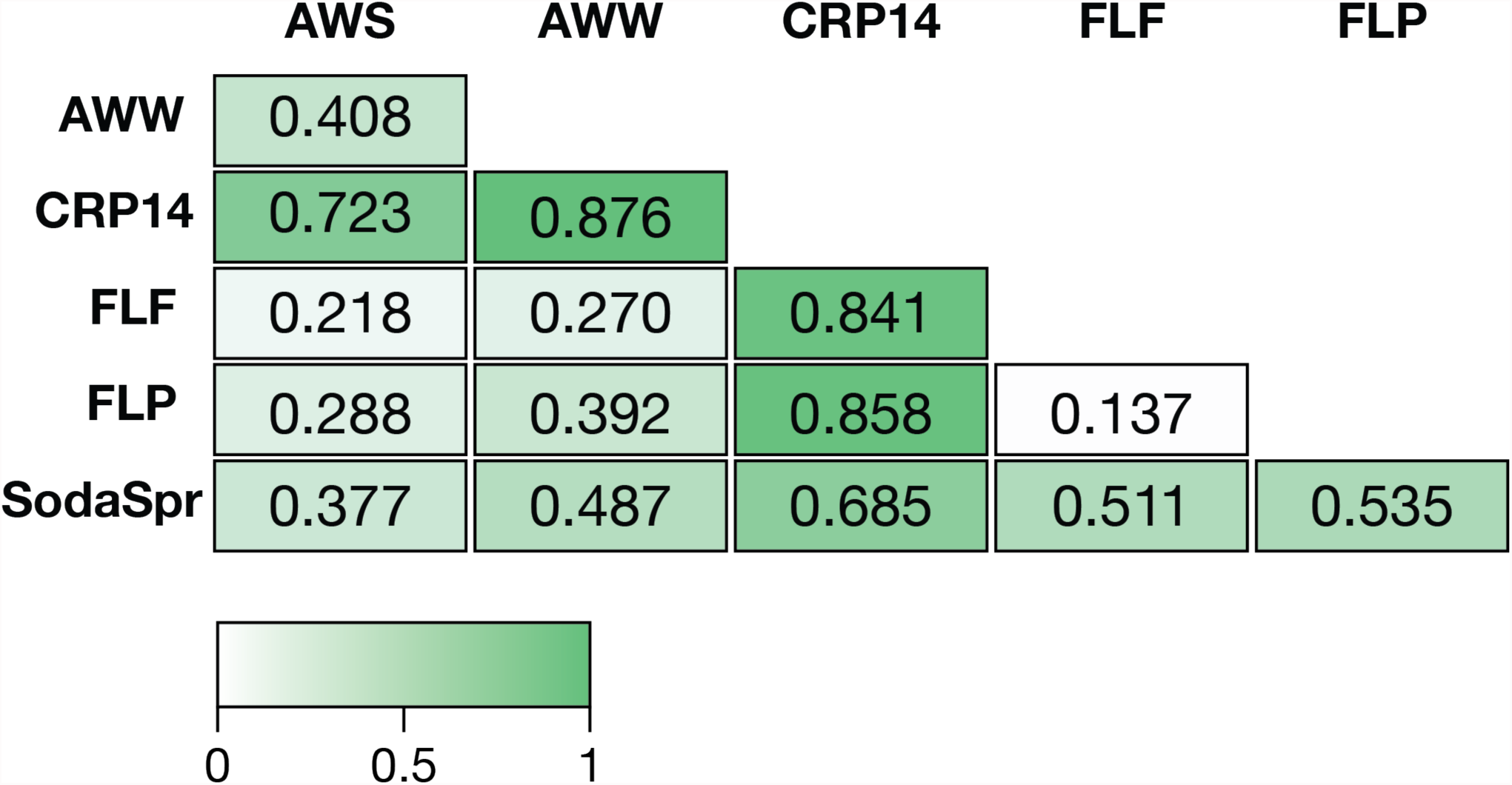
Analysis of similarity (ANOSIM) R-values showing differences in microbial community composition between each of the six substrate arrays.

Inspection of the most abundant organisms in each substrate arrays reveals that many of the enrichments in each substrate array are inhabited by one or several dominant taxa (Figure 5). These taxa, of which OTUs classified as *Pseudomonas*, *Ralstonia*, the family (fa.) *Enterobacteriaceae*, *Burkholderia*, and *Agrobacterium* are the most abundant, are not the most abundant taxa in the inocula. Sequences classified in the genus *Pseudomonas* account for 36% of all sequences across all 6 substrate arrays, but have an average abundance of only 0.26±0.17% in the inocula. In five of the six substrate arrays (except CRP14), the relative abundance of *Pseudomonas* is on average 61–300 fold greater relative to the inoculum. In CRP14 *Pseudomonas* was only enriched four-fold, on average, and comprised < 0.3% of the total diversity seen across all wells and was of > 1% abundance in only 3 wells (3-11% in those wells). Others, like *Ralstonia*, fa. *Enterobacteriaceae*, *Burkholderia*, and *Agrobacterium* are considerably enriched in many if not most of the wells within individual substrate arrays, but are largely or entirely absent in others (Figure 5).

**Figure 5.**
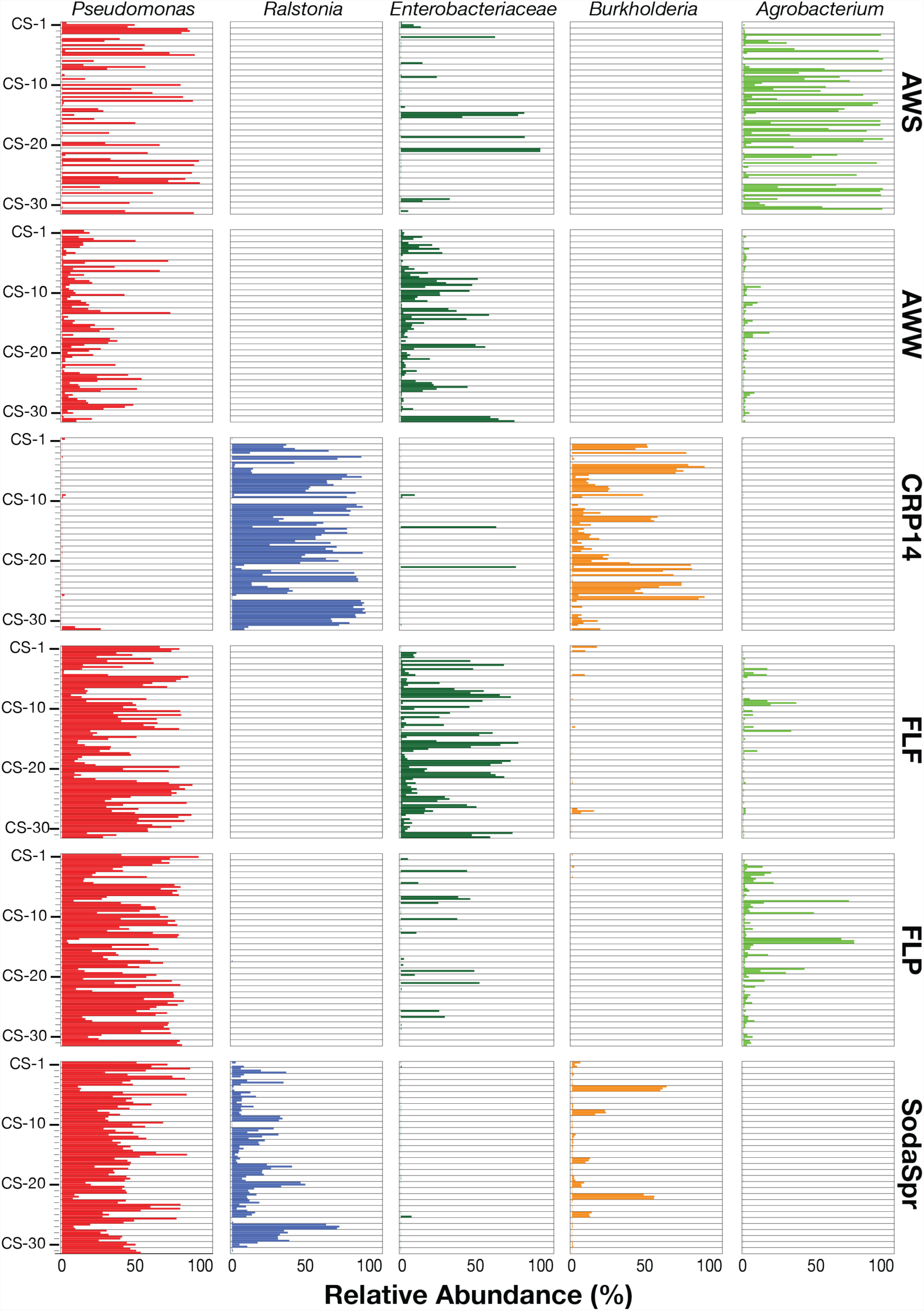
Relative abundance of the top five taxa (*Pseudomonas*, *Ralstonia*, *Agrobacterium*, *Burkholderia*, and *Enterobacteriaceae*) in the EcoPlate enrichments, with triplicate samples grouped by carbon amendment.

This pattern is also true for less broadly abundant taxa like *Comamonas* and the fa. *Xanthomonadaceae*. The distribution of all OTUs enriched or depleted (based on the edgeR metric) in the substrate arrays compared to the inocula is shown in Figure 6. Overall, these data confirm the observational data shown in Figure 5, that the OTUs most abundant in the enrichments are largely members of the *Proteobacteria* (*Pseudomonas*, *Ralstonia*, *Agrobacterium*) and the *Bacteroidetes* (*Chryseobacterium*, *Flavobacterium*, *Sphingobacterium*, *Pedobacter*, and *Wautersiella*). OTUs classified as other phyla (e.g., *Actinobacteria*, *Verrucomicrobia*, and *Acidobacteria*) are considerably less abundant in the enrichments than they are in the original environments. The distribution of specific OTUs also varies within taxonomic groups. As seen in Figure 7, in four of the five substrate arrays where *Pseudomonas* is abundant (AWS, AWW, FLF, and FLP), a single OTU predominates (labelled OTU A). While OTU A is present in some enrichments inoculated with material from the SodaSpr site, a different OTU (OTU H) is much more dominant. Overall, of the 10 taxonomic groups that were found to be above 1% average abundance in the substrate arrays (*Pseudomonas*, *Ralstonia*, fa. *Enterobacteriaceae*, *Agrobacterium*, *Burkholderia*, fa. *Aeromonadaceae*, *Chryseobacterium*, fa. *Xanthomonadaceae*, *Delftia*, *Comamonas*), sequences from all these together comprise only 1.1-2.5% of the total diversity of the inocula in the five soil arrays and 8.0% of the diversity in the wetland water array.

**Figure 6.**
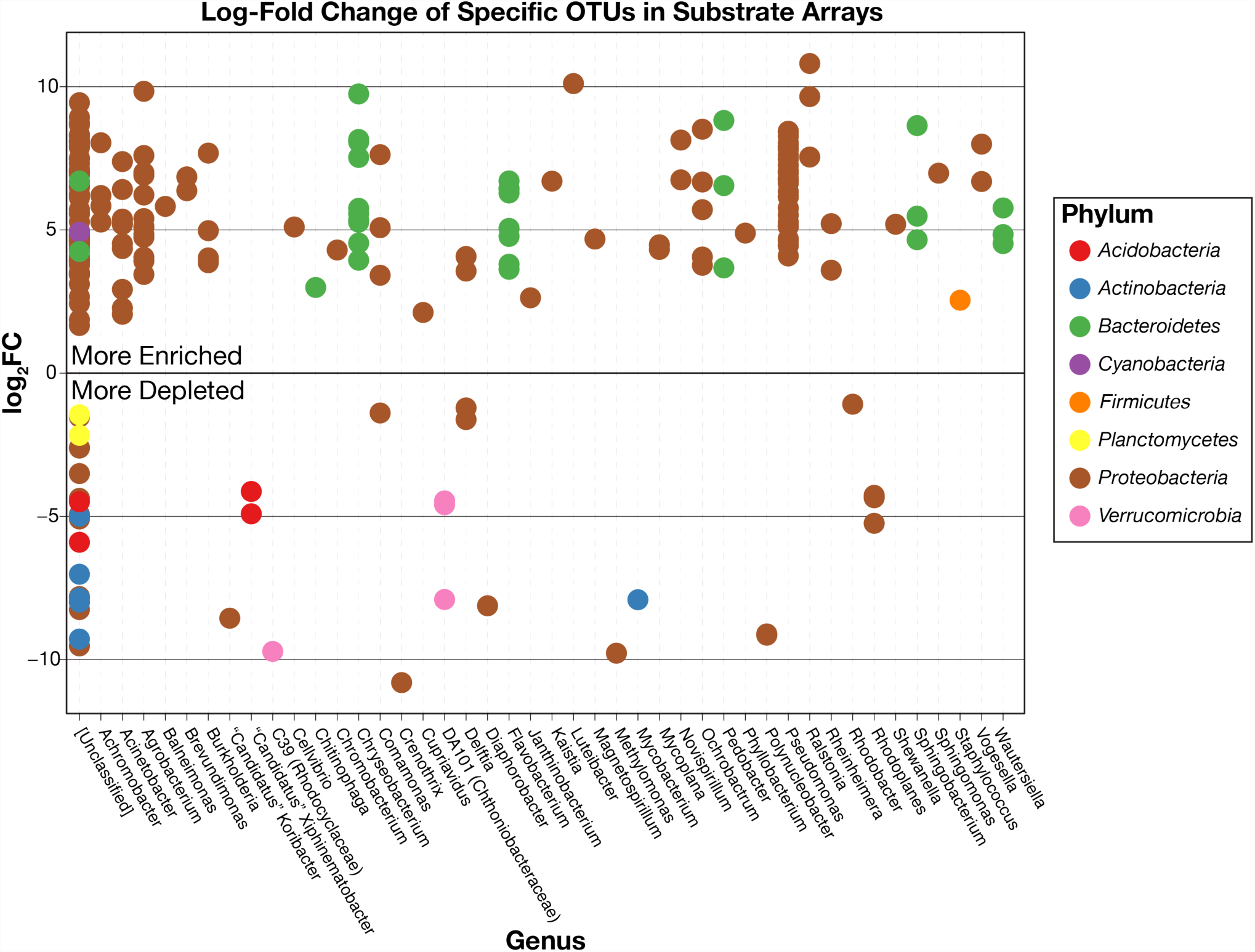
Log-fold (base two) change in the relative abundance of individual OTUs in the substrate array enrichments relative to the original inoculum based on the edgeR package. Positively enriched OTUs are more abundant, on average, in the substrate arrays than in the inocula while negatively enriched OTUs are more abundant in the inocula. OTUs are separated horizontally by genus and colored by phylum.

**Figure 7.**
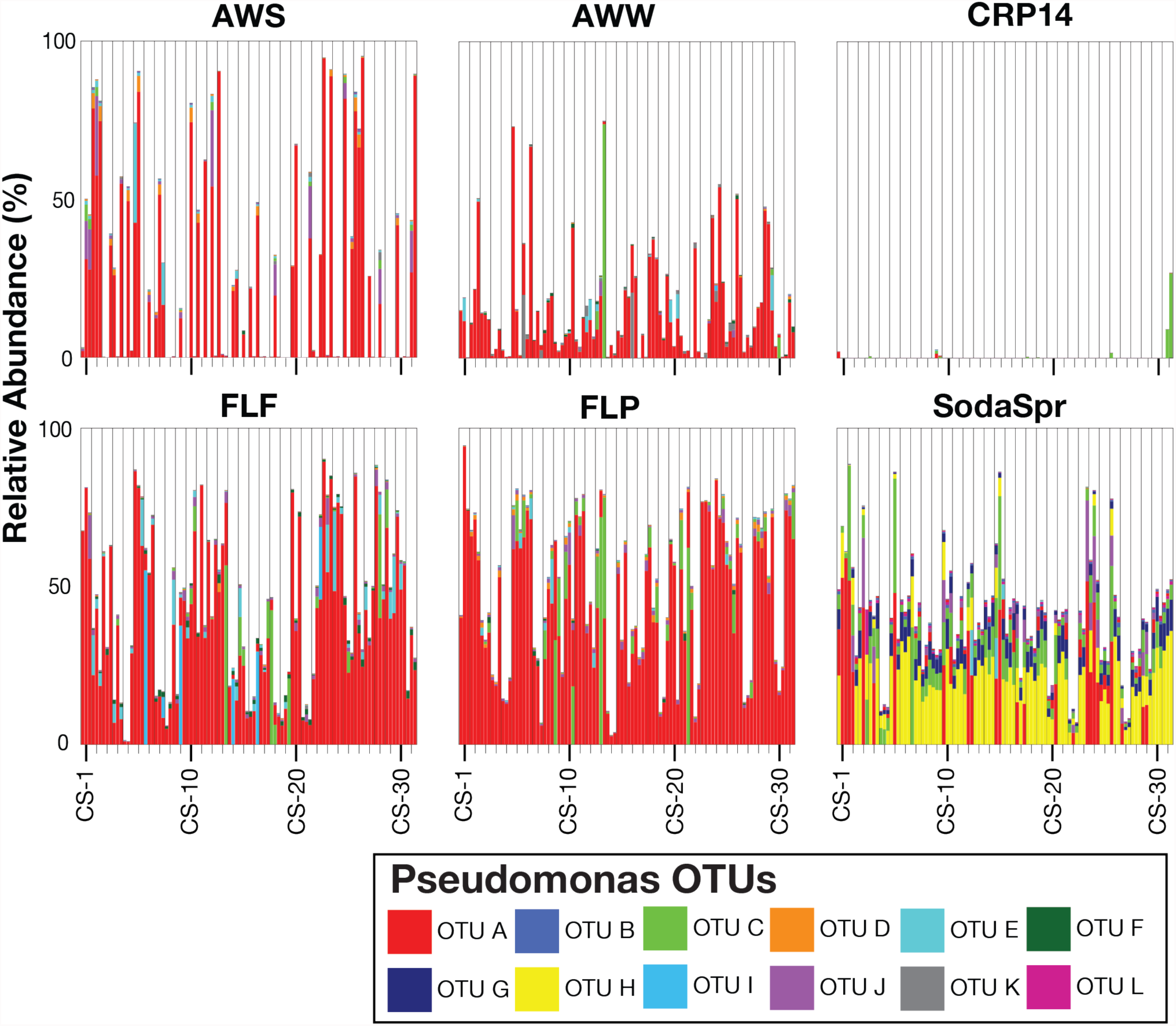
Relative abundance of individual OTUs classified within the genus *Pseudomonas*.

Additional pairwise ANOSIM calculations were conducted to test for systematic differences in community composition between enrichments amended with a specific substrate category (amines, amino acids, carbohydrates, polymers, and phenolic compounds) and within each individual substrate itself. This test compared whether communities from across all six substrate arrays amended with a specific class of carbon compound (e.g., amines) were, on average, more similar to each other than to communities amended with a different class of compounds (e.g., carbohydrates). While no statistically significant correlations held observed across all six substrate arrays, several statistically significant ANOSIM values were observed for several substrate categories within a given environment. Most significantly, comparing enrichments amended with a carbohydrate with those amended with a phenolic compound yield statistically significant ANOSIM values (R_ANOSIM_ = 0.483-0.597) in 3 of the 6 environmental enrichments: wetland water, temperate forest soil, and subalpine forest soil. None of the other within-environment comparisons yielded significant results for more than 1 of the 6 environmental sources.

Additional comparisons between phenolic compound-amended and carbohydrate-amended enrichments made using edgeR found that 42 OTUs above the chosen variance of 10^−5^ total threshold (McMurdie and Holmes, 2013) were differentially abundant (Figure 8). Overall, we found that OTUs within the phylum *Bacteroidetes* were generally more abundant in the phenolic-amended enrichments, while OTUs classified within the *Proteobacteria* were generally more abundant in carbohydrate-amended enrichments. OTUs classified in the genera *Wautersiella*, *Comamonas*, *Sphingobacterium*, *Chitinophaga*, *Sphingobium*, and *Staphylococcus* were more abundant in the phenolic-amended microcosms.

**Figure 8.**
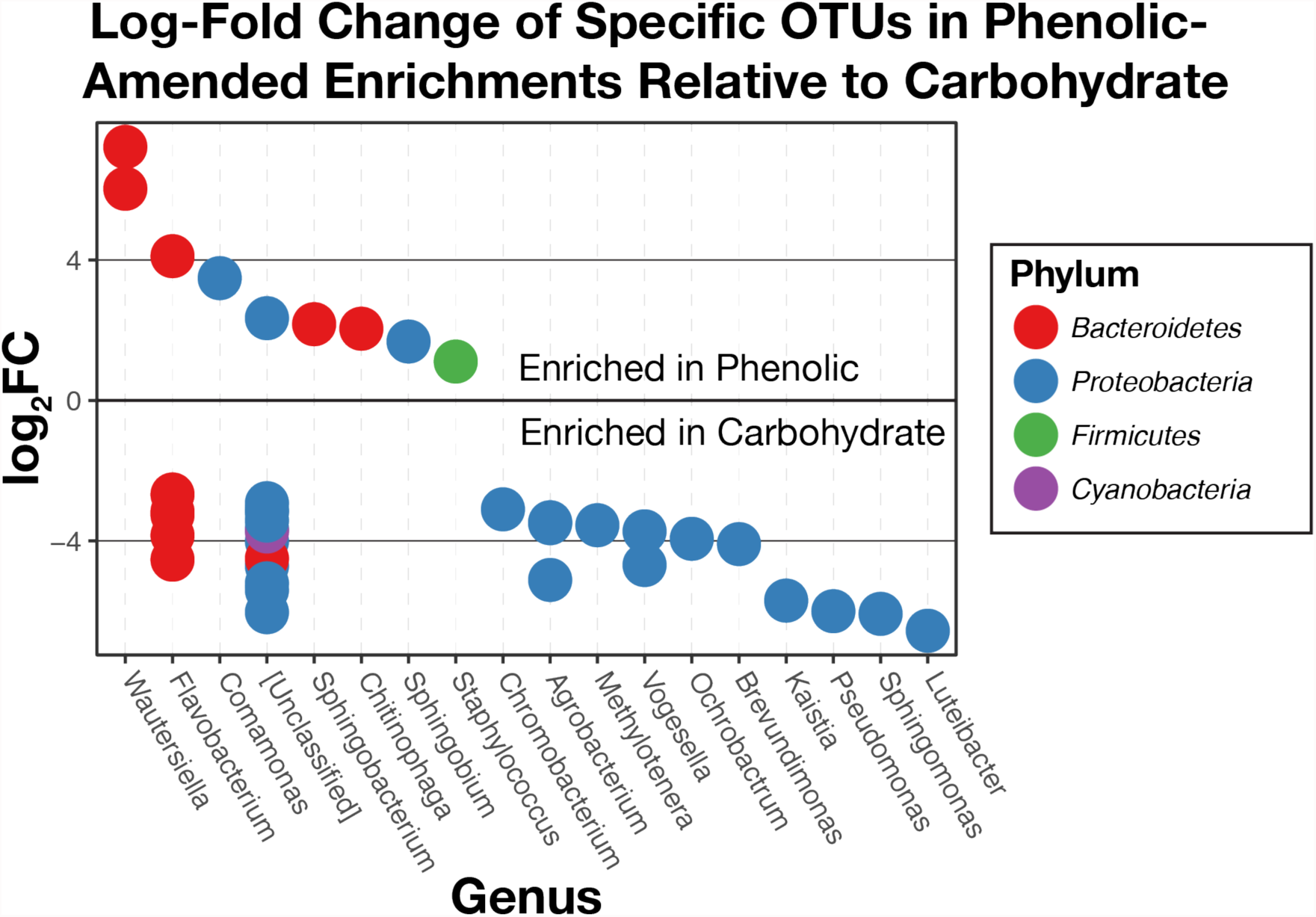
Log2-fold changes of differentially abundant operational taxonomic units (OTUs) in substrate arrays wells given phenolic substrates compared to carbohydrates. OTUs that are positively enriched are more abundant in phenolic enrichments. Those that are negatively enriched are more abundant in carbohydrate enrichments.

## 4 Discussion

The distinct communities of microorganisms that grow to dominate each enrichment within the substrate arrays bear the strong influence of the initial composition of the soil or water used as inoculum. As can be seen from the NMDS plot in Figure 3, the source of inoculum is by far the strongest signal in differentiating the microbial communities in these substrate arrays. The statistical significance of this result is confirmed by ANOSIM based on weighted UniFrac^1^, where the resulting R-values recapitulate the trends observed in the NMDS plot (Figure 4). Furthermore, these results show that, for the most part, the communities in the substrate arrays from the most divergent inocula [tropical (CRP14) and subalpine (SodaSpr) forest soil] remain divergent from those sourced from temperate soil inocula (AWS, FLP, FLF), which beta diversity comparisons indicate are more similar to each other (Figure S2). The exception is the wetland surface water (AWW), which Figure S2 shows has the most divergent community composition of the six inocula prior to enrichment. Following the 72-hour incubation, however, the communities that arise in the substrate arrays inoculated with AWW material appear much more similar to the arrays inoculated with temperate soil than to those inoculated with soil from CRP14 or SodaSpr (Figure 3). This suggests that the generalists like *Pseudomonas*, fa. *Enterobacteriaceae*, and *Agrobacterium* that grow to dominate the substrate arrays share a similar baseline activity across all the temperate environments and are therefore able to be enriched to a similar degree in the arrays. Still, the smaller but still significant R_ANOSIM_ values between some of the terrestrial enrichments (e.g., AWW and AWS) indicate the existence of smaller but significant differences between these environments.

Despite the strong influence of the initial inoculum, the overall structure of the microbial communities that grow to inhabit these substrate arrays bears little resemblance to that of the parent material, as measured by both alpha (Figure 2) and beta diversity (Figure S1). By and large, the enrichments in each substrate array are primarily dominated by one or several OTUs most closely related to aerobic, heterotrophic generalists like *Pseudomonas*, *Burkholderia*, *Ralstonia*, fa. *Enterobacteriaceae*, and others (Figure 5, Figure 6). *Pseudomonas* in particular dominated five of the six substrate arrays, although it was nearly absent from enrichments in the array inoculated with soil from site CRP14. *Pseudomonas* and its relatives are well-known as metabolic generalists and are frequently found in aerobic soil environments. Indeed, in their pre-phylogenetic taxonomic classification, “pseudomonads” were defined largely by “their most striking and ecologically significant group character… namely, the ability to use a wide variety of organic compounds as carbon and energy sources for aerobic growth” (Stanier et al., 1966). Numerous microcosm and enrichment studies from soils under aerobic conditions have found abundant growth of *Pseudomonas* (Greene et al., 2000; Eriksson et al., 2003; Ma et al., 2006), although these are often limited in scope to a few replicates or the utilization of a particular compound of interest.

An exception to this repeated occurrence of *Pseudomonas* was in the apparent swapping of *Pseudomonas* for *Burkholderia* and *Ralstonia* across the CRP14-inoculated enrichments. Like *Pseudomonas*, *Burkholderia* is a metabolic generalist well-distributed in soil environments (Cho and Tiedje, 2000; Salles et al., 2004; Janssen, 2006; Compant et al., 2008; Lauber et al., 2009; Stopnisek et al., 2014). *Ralstonia* is also commonly genera found in soil environments (Janssen, 2006), although most studies of this genus focus on its role as soilborne plant pathogen (Castillo and Greenberg, 2007; Leonard et al., 2017) with a wide distribution in warm and tropical climates (Genin and Boucher, 2004). Both *Burkholderia* and *Ralstonia* share many characteristics with *Pseudomonas*; both were at one time themselves classified as part of that genus (Yabuuchi et al., 1992; Yabuuchi et al., 1995) although they are now recognized as *Betaproteobacteria* while *Pseudomonas* is classified as a member of the *Gammaproteobacteria*. The abundance of these two genera in the CRP14- amended substrate array suggests an optimization of the community from the tropical forest system to the set of niches found in the source environment to the exclusion of *Pseudomonas*. There is likely some consistent factor associated with those niches, however, so that a “distinct generalist” is enriched consistently over time in one environment but not another.

When comparing the substrate array enrichments by the chemical class of substrate (amines, amino acids, carbohydrates, polymers, or phenolic compounds) rather than source environment, we found that significant differences in community composition exist primarily between those wells given a carbohydrate as a carbon source and those enriched with a phenolic compound (Figure 8). These differences were observed across three of the six environmental inocula: AWW, FLF, and SodaSpr, while none of the other pairwise comparisons between carbon source class showed a significant difference in more than one substrate array. Taken together, these results suggest that the co-occurrence of minimal community members within these three environments and grown on carbohydrates or phenolics are likely non-random associations. Like the most abundant taxa across the substrate arrays, most of the differentially-abundant OTUs (Figure 8) between phenolic- and carbohydrate-amended enrichments are most closely related to aerobic heterotrophs. Interestingly, the closest neighbors of several of the OTUs that are differentially abundant in phenolic-amended enrichments are organisms that are specifically known for degrading aromatic compounds. For example, members of the genus *Comamonas* have previously been isolated and studied based on their ability to degrade polycyclic aromatic hydrocarbons (Goyal and Zylstra, 1996) while isolated members of the genus *Sphingobium* are known for the ability to degrade phenolic compounds such as nonylphenols (Ushiba et al., 2003) and chlorophenols (Cai and Xun, 2002) as well as other aromatic hydrocarbons (Cunliffe and Kertesz, 2006). Carbohydrate-amended enrichments were more enriched OTUs classified as *Luteibacter*, *Sphingomonas*, *Pseudomonas*, *Kaistia*, *Brevundimonas*, *Ochrobactrum*, *Vogesella*, and others. Unsurprisingly, known isolates from these groups are primarily aerobic heterotrophs with wide metabolic diversity, including the ability to grow on carbohydrates as an energy source (Lessie and Phibbs, 1984; Lebuhn et al., 2000; Im et al., 2004; de Boer et al., 2007) and are commonly found in a variety of terrestrial environments (Fredrickson et al., 1999; Cho and Tiedje, 2000; Spiers et al., 2000; Lauber et al., 2009). Despite being more enriched in carbohydrate-amended wells, *Sphingomonas* spp. have also shown the ability to grow on aromatic compounds (Fredrickson et al., 1995).

Taken together, these findings provide substantial insight into the forces that shape the composition of the microbiome. Microbial communities, like all communities of living organisms, are thought to be shaped by two types of ecological factors: those that are relatively deterministic are dubbed “niche” while those that are more stochastic are “neutral” (Stegen et al., 2012). Niche factors that influence microbial community dynamics include environmental factors such as temperature or pH, the presence or absence of a particular nutrient (e.g. carbon, nitrogen, phosphorus, etc.) or the physical structure of the environment itself (e.g. the porosity or mineralogy of the soil matrix). Neutral forces, conversely, include random birth/death events, migrations of a particular population from one area to another, and other probabilistic events (Chase and Myers, 2011; Stegen et al., 2013). Evidence of the significance of both processes has been observed throughout a variety of environments (Gilbert et al., 2012; Shade et al., 2012; Handley et al., 2014). This poses a significant challenge to creating phylogenetically-resolved predictive models of microbial activity (Graham et al., 2016), particularly considering the breathtaking diversity of soil microbial communities (Howe et al., 2014).

In the minimized communities that inhabit our substrate arrays, we see evidence of both niche and neutral processes shaping the overall community composition. Niche selection clearly determines the overall trajectory of the response of the initial inoculum to changes in environmental conditions, as the organisms that grow to dominate the wells of the substrate arrays are aerobic heterotrophs, matching the oxic and carbon-rich conditions found in the wells. Further niche selection appears evident in the distinction between carbohydrate- and phenolic-amended enrichments, as specific populations closely related to isolates known for degrading aromatic compounds become significantly more abundant in the presence of phenolic compounds (Figure 8). Still, despite the fact that the source of inoculum is a significant determinant of the overall community structure (Figure 3), probabilistic events still appear to control some of the finer aspects of the composition of the substrate array communities. For example, the extent to which triplicate wells on the same substrate array ended up with the same community composition varied depending on both the substrate and the population. For example, when populations classified as *Burkholderia* became abundant in wells inoculated with soil from the SodaSpr site, the abundance was observed uniformly across all three replicates (Figure 6). Conversely for some of carbon sources in wells amended with AWS soil, two of the three triplicates will contain >90% *Pseudomonas* while the third well will be similarly dominated by sequences classified as fa. *Enterobacteriaceae* or *Agrobacterium* (Figure 6). Because the inocula were rigorously diluted, dispersed, and homogenized prior to being added to the substrate arrays, it appears unlikely that this is an artifact of sample preparation. Instead, stochasticity and imperceptible differences in the initial inoculum likely play at least some role in determining the overall trajectory of change in community composition. This has been seen elsewhere, as in Kwon et al.’s (2016) observation of two distinct mineralogical and microbiological trajectories in replicate microcosms amended with glucose and ferric iron.

The parallelized enrichment strategy we employ here provides an initial framework to capture some of the underlying and potentially novel interactions between microbial community members found in soil and water in a high-throughput manner. This work also has the potential to inform the intentional design of microbial communities (synthetic ecology). Current work in our laboratory is focused on testing the stability of these “minimal communities” in continuous culture systems with the hope of creating hybrid synthetic communities that are sourced directly from specific environments but streamlined in the laboratory. The isolation of a consistently reoccurring generalist provides an anchor within the community for genetic engineering. A variety of genetic tools are available for *Pseudomonas* and other taxa seen here, providing a potential platform for customization of minimal communities for industrial and biomedical applications. Moving forward, as desire increases to engineer microbial communities either by strain selection or outright genetic manipulation, the use of highly parallelized microbial enrichments promises to provide novel insights that will allow us to predict with greater accuracy the interactions that contribute to emergent properties of microbial communities, such as community stability.

## 5 Conflict of Interest

The authors declare that the research was conducted in the absence of any commercial or financial relationships that could be construed as a potential conflict of interest.

## 6 Author Contributions

DA, TF, and KK designed the project. TF collected samples and conducted the experiments. JK extracted DNA and prepared sequencing libraries. SG and SO carried out DNA sequencing. TF and DA analyzed the data and wrote the manuscript. All authors reviewed and approved the final manuscript.

## 7 Funding

This research was funded by Argonne National Laboratory, U.S. Department of Energy (DOE) Office of Science, under DOE contract DE-AC02-06CH11357.

## 8 Acknowledgments

The authors thank T. Beemer for his assistance with wetland classification. We also thank S. O’Brien, E. O’Loughlin, S. Alvarez-Clare, and P. Weisenhorn, for helpful discussions and insights that substantially improved this manuscript.

## Footnotes

1 ANOSIM was also performed using the Bray-Curtis similarity coefficient and exhibited similar trends (data not shown).

## 13 Supplementary Material

**Figure S1.**
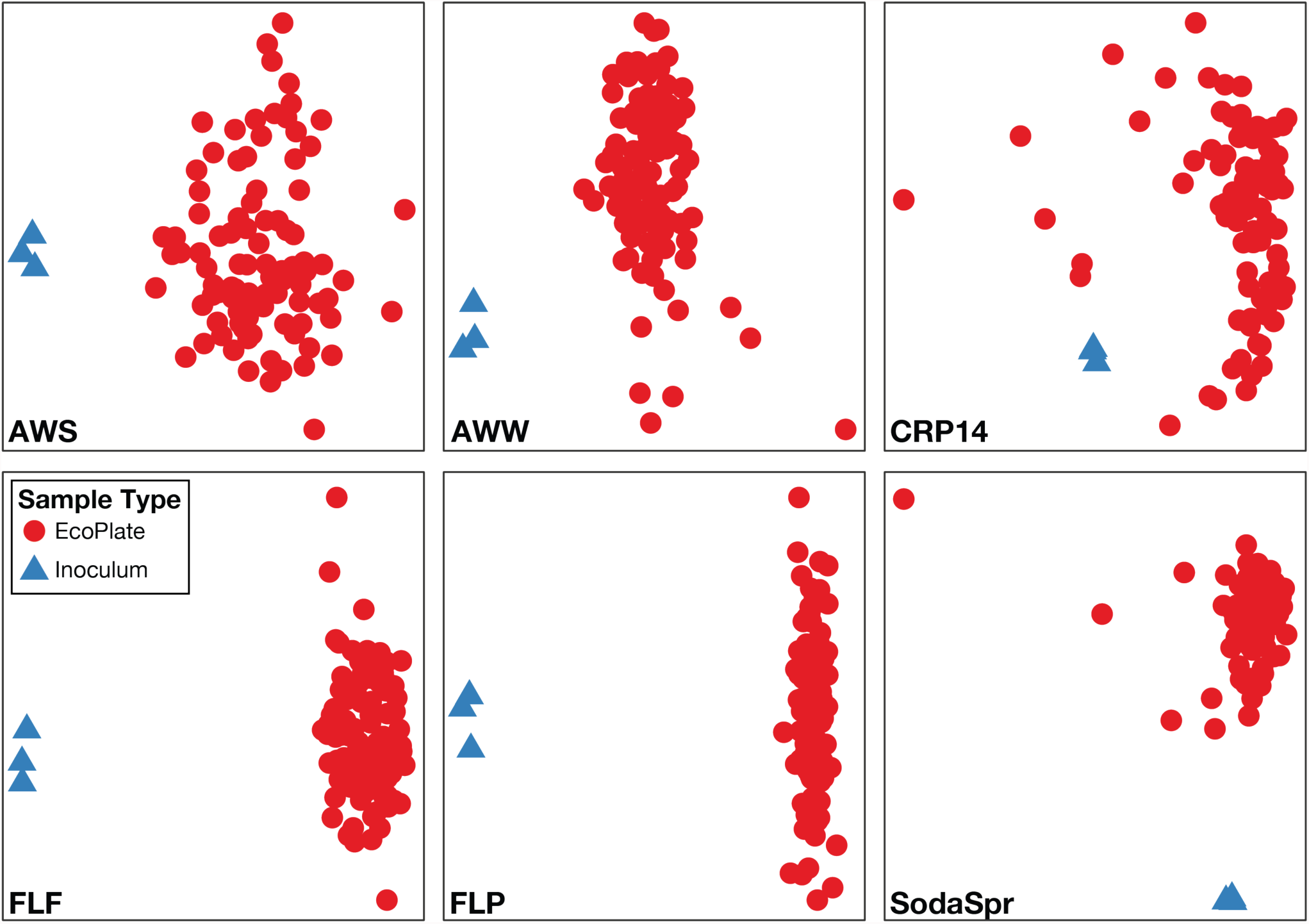
NMDS plots highlighting the difference in microbial community structure between environmental inocula (blue) and EcoPlate enrichments (red). Each individual plot includes all samples from a particular environment: freshwater wetland soil (AWS), freshwater wetland water (AWW), tropical forest soil (CRP14), temperate forest soil (FLF), temperate prairie soil (FLF), subalpine forest soil (SodaSpr).

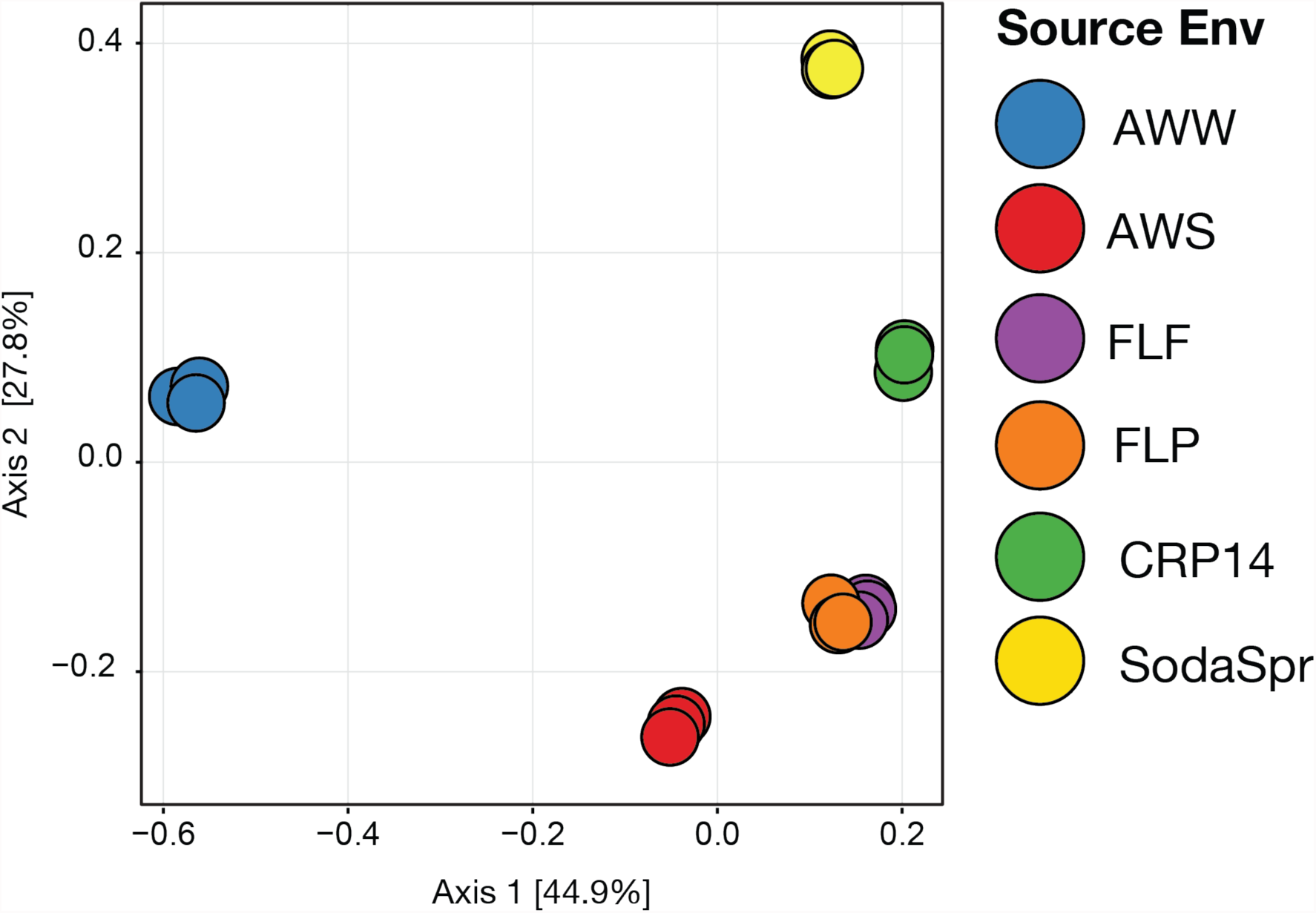
**Figure S2**. Principal coordinates (PCoA) plot of the differences between triplicate samples of the environmental inocula used in substrate array experiments: freshwater wetland soil (AWS), freshwater wetland water (AWW), tropical forest soil (CRP14), temperate forest soil (FLF), temperate prairie soil (FLF), subalpine forest soil (SodaSpr). Primary source of variation along Axis 1 is between soil communities and planktonic cells from wetland surface water (AWW). Additional variation along axis two appears to differentiate between microbial communities in Midwestern soil (AWS, FLF, and FLP) compared to subalpine (SodaSpr) and tropical (CRP14) soil.

## 9 References

Alvarez-Clare, S., Mack, M.C., and Brooks, M. (2013). A direct test of nitrogen and phosphorus limitation to net primary productivity in a lowland tropical wet forest. Ecology 94, 1540-1551.

Bailey, V.L., Smith, A.P., Tfaily, M., Fansler, S.J., and Bond-Lamberty, B. (2017). Differences in soluble organic carbon chemistry in pore waters sampled from different pore size domains. Soil Biology and Biochemistry 107, 133-143.

Baldwin, D.S., Rees, G.N., Mitchell, A.M., Watson, G., and Williams, J. (2006). The short-term effects of salinization on anaerobic nutrient cycling and microbial community structure in sediment from a freshwater wetland. Wetlands 26, 455-464.

Bartscht, K., Cypionka, H., and Overmann, J. (1999). Evaluation of cell activity and of methods for the cultivation of bacteria from a natural lake community. FEMS Microbiology Ecology 28, 249-259.

Bates, S.T., Berg-Lyons, D., Caporaso, J.G., Walters, W.A., Knight, R., and Fierer, N. (2011). Examining the global distribution of dominant archaeal populations in soil. The ISME Journal 5, 908-917.

Bochner, B.R. (1989). Sleuthing out bacterial identities. Nature 339, 157-158.

Bochner, B.R., Gadzinski, P., and Panomitros, E. (2001). Phenotype microarrays for high-throughput phenotypic testing and assay of gene function. Genome Research 11, 1246-1255.

Brennan, R.A., Sanford, R.A., and Werth, C.J. (2006). Chitin and corncobs as electron donor sources for the reductive dechlorination of tetrachloroethene. Water Research 40, 2125-2134.

Cai, M., and Xun, L. (2002). Organization and regulation of pentachlorophenol-degrading genes in *Sphingobium chlorophenolicum* ATCC 39723. Journal of Bacteriology 184, 4672-4680.

Caporaso, J.G., Kuczynski, J., Stombaugh, J., Bittinger, K., Bushman, F.D., Costello, E.K., Fierer, N., Pena, A.G., Goodrich, J.K., Gordon, J.I., Huttley, G.A., Kelley, S.T., Knights, D., Koenig, J.E., Ley, R.E., Lozupone, C.A., McDonald, D., Muegge, B.D., Pirrung, M., Reeder, J., Sevinsky, J.R., Turnbaugh, P.J., Walters, W.A., Widmann, J., Yatsunenko, T., Zaneveld, J., and Knight, R. (2010). QIIME allows analysis of high-throughput community sequencing data. Nature Methods 7, 335-336.

Caporaso, J.G., Lauber, C.L., Walters, W.A., Berg-Lyons, D., Huntley, J., Fierer, N., Owens, S.M., Betley, J., Fraser, L., Bauer, M., Gormley, N., Gilbert, J.A., Smith, G., and Knight, R. (2012). Ultra-high-throughput microbial community analysis on the Illumina HiSeq and MiSeq platforms. The ISME Journal 6, 1621-1624.

Caporaso, J.G., Lauber, C.L., Walters, W.A., Berg-Lyons, D., Lozupone, C.A., Turnbaugh, P.J., Fierer, N., and Knight, R. (2011). Global patterns of 16S rRNA diversity at a depth of millions of sequences per sample. Proceedings of the National Academy of Sciences 108, 4516-4522.

Castillo, J.A., and Greenberg, J.T. (2007). Evolutionary dynamics of *Ralstonia solanacearum*. Applied and Environmental Microbiology 73, 1225-1238.

Chase, J.M., and Myers, J.A. (2011). Disentangling the importance of ecological niches from stochastic processes across scales. Philosophical Transactions of the Royal Society B: Biological Sciences 366, 2351-2363.

Cho, J.-C., and Tiedje, J.M. (2000). Biogeography and degree of endemicity of fluorescent Pseudomonas strains in soil. Applied and Environmental Microbiology 66, 5448-5456.

Clarke, K.R., and Warwick, R.M. (2001). Change in Marine Communities: An Approach to Statistical Analysis and Interpretation. Plymouth, UK: PRIMER-E, Ltd.

Compant, S., Nowak, J., Coenye, T., Clément, C., and Ait Barka, E. (2008). Diversity and occurrence of *Burkholderia* spp. in the natural environment. FEMS Microbiology Reviews 32, 607-626.

Cunliffe, M., and Kertesz, M.A. (2006). Effect of *Sphingobium yanoikuyae* B1 inoculation on bacterial community dynamics and polycyclic aromatic hydrocarbon degradation in aged and freshly PAH-contaminated soils. Environmental Pollution 144, 228-237.

Dalcin Martins, P., Hoyt, D.W., Bansal, S., Mills, C.T., Tfaily, M., Tangen, B.A., Finocchiaro, R.G., Johnston, M.D., McAdams, B.C., Solensky, M.J., Smith, G.J., Chin, Y.-P., and Wilkins, M.J. (2017). Abundant carbon substrates drive extremely high sulfate reduction rates and methane fluxes in Prairie Pothole Wetlands. Global Change Biology.

de Boer, W., Wagenaar, A.-M., Klein Gunnewiek, P.J.A., and van Veen, J.A. (2007). In vitro suppression of fungi caused by combinations of apparently non-antagonistic soil bacteria. FEMS Microbiology Ecology 59, 177-185.

Eren, A.M., Maignien, L., Sul, W.J., Murphy, L.G., Grim, S.L., Morrison, H.G., and Sogin, M.L. (2013). Oligotyping: differentiating between closely related microbial taxa using 16S rRNA gene data. Methods in Ecology and Evolution 4, 1111-1119.

Eriksson, M., Sodersten, E., Yu, Z., Dalhammar, G., and Mohn, W.W. (2003). Degradation of polycyclic aromatic hydrocarbons at low temperature under aerobic and nitrate-reducing conditions in enrichment cultures from northern soils. Applied and Environmental Microbiology 69, 275-284.

Flynn, T.M., Sanford, R.A., Ryu, H., Bethke, C.M., Levine, A.D., Ashbolt, N.J., and Santo Domingo, J.W. (2013). Functional microbial diversity explains groundwater chemistry in a pristine aquifer. BMC Microbiology 13, 146.

Fredrickson, J.K., Balkwill, D.L., Drake, G.R., Romine, M.F., Ringelberg, D.B., and White, D.C. (1995). Aromatic-degrading *Sphingomonas* isolates from the deep subsurface. Applied and Environmental Microbiology 61, 1917-1922.

Fredrickson, J.K., Balkwill, D.L., Romine, M.F., and Shi, T. (1999). Ecology, physiology, and phylogeny of deep subsurface *Sphingomonas* sp. Journal of Industrial Microbiology and Biotechnology 23, 273-283.

Genin, S., and Boucher, C. (2004). Lessons learned from the genome analysis of *Ralstonia solanacearum*. Annual Review of Phytopathology 42, 107-134.

Gilbert, J.A., Steele, J.A., Caporaso, J.G., Steinbruck, L., Reeder, J., Temperton, B., Huse, S., McHardy, A.C., Knight, R., Joint, I., Somerfield, P., Fuhrman, J.A., and Field, D. (2012). Defining seasonal marine microbial community dynamics. The ISME Journal 6, 298-308.

Goyal, A.K., and Zylstra, G.J. (1996). Molecular cloning of novel genes for polycyclic aromatic hydrocarbon degradation from *Comamonas testosteroni* GZ39. Applied and Environmental Microbiology 62, 230-236.

Graham, E.B., Knelman, J.E., Schindlbacher, A., Siciliano, S., Breulmann, M., Yannarell, A., Beman, J.M., Abell, G., Philippot, L., Prosser, J., Foulquier, A., Yuste, J.C., Glanville, H.C., Jones, D.L., Angel, R., Salminen, J., Newton, R.J., Bürgmann, H., Ingram, L.J., Hamer, U., Siljanen, H.M.P., Peltoniemi, K., Potthast, K., Bañeras, L., Hartmann, M., Banerjee, S., Yu, R.-Q., Nogaro, G., Richter, A., Koranda, M., Castle, S.C., Goberna, M., Song, B., Chatterjee, A., Nunes, O.C., Lopes, A.R., Cao, Y., Kaisermann, A., Hallin, S., Strickland, M.S., Garcia-Pausas, J., Barba, J., Kang, H., Isobe, K., Papaspyrou, S., Pastorelli, R., Lagomarsino, A., Lindström, E. S., Basiliko, N., and Nemergut, D.R. (2016). Microbes as engines of ecosystem function: When does community structure enhance predictions of ecosystem processes? Frontiers in Microbiology 7.

Greene, E.A., Kay, J.G., Jaber, K., Stehmeier, L.G., and Voordouw, G. (2000). Composition of soil microbial communities enriched on a mixture of aromatic hydrocarbons. Applied and Environmental Microbiology 66, 5282-5289.

Gryta, A., Frąc, M., and Oszust, K. (2014). The application of the Biolog EcoPlate approach in ecotoxicological evaluation of dairy sewage sludge. Applied Biochemistry and Biotechnology 174, 1434-1443.

Haack, S.K., Fogarty, L.R., West, T.G., Alm, E.W., McGuire, J.T., Long, D.T., Hyndman, D.W., and Forney, L.J. (2004). Spatial and temporal changes in microbial community structure associated with recharge-influenced chemical gradients in a contaminated aquifer. Environmental Microbiology 6, 438-448.

Handley, K.M., Wrighton, K.C., Miller, C.S., Wilkins, M.J., Kantor, R.S., Thomas, B.C., Williams, K.H., Gilbert, J.A., Long, P.E., and Banfield, J.F. (2014). Disturbed subsurface microbial communities follow equivalent trajectories despite different structural starting points. Environmental Microbiology.

Hitzl, W., Rangger, A., Sharma, S., and Insam, H. (1997). Separation power of the 95 substrates of the BIOLOG system determined in various soils. FEMS Microbiology Ecology 22, 167-174.

Howe, A.C., Jansson, J.K., Malfatti, S.A., Tringe, S.G., Tiedje, J.M., and Brown, C.T. (2014). Tackling soil diversity with the assembly of large, complex metagenomes. Proceedings of the National Academy of Sciences 111, 4904-4909.

Hug, L.A., Thomas, B.C., Brown, C.T., Frischkorn, K.R., Williams, K.H., Tringe, S.G., and Banfield, J.F. (2015). Aquifer environment selects for microbial species cohorts in sediment and groundwater. The ISME Journal 9, 1846-1856.

Im, W.-T., Yokota, A., Kim, M.-K., and Lee, S.-T. (2004). *Kaistia adipata* gen. nov., sp. nov., a novel α-proteobacterium. The Journal of General and Applied Microbiology 50, 249-254.

Janssen, P.H. (2006). Identifying the dominant soil bacterial taxa in libraries of 16S rRNA and 16S rRNA genes. Applied and Environmental Microbiology 72, 1719-1728.

King, G.M. (2003). Contributions of atmospheric CO and hydrogen uptake to microbial dynamics on recent Hawaiian volcanic deposits. Applied and Environmental Microbiology 69, 4067-4075.

Kirk, M.F., Santillan, E.F.U., Sanford, R.A., and Altman, S.J. (2013). CO_2_-induced shift in microbial activity affects carbon trapping and water quality in anoxic bioreactors. Geochimica et Cosmochimica Acta 122, 198-208.

Kirk, M.F., Wilson, B.H., Marquart, K.A., Zeglin, L.H., Vinson, D.S., and Flynn, T.M. (2015). Solute concentrations influence microbial methanogenesis in coal-bearing strata of the Cherokee basin, USA. Frontiers in Microbiology 6.

Konopka, A., Oliver, L., and Turco, J.R.F. (1998). The use of carbon substrate utilization patterns in environmental and ecological microbiology. Microbial Ecology 35, 103-115.

Kraft, B., Tegetmeyer, H.E., Meier, D., Geelhoed, J.S., and Strous, M. (2014). Rapid succession of uncultured marine bacterial and archaeal populations in a denitrifying continuous culture. Environmental Microbiology 16, 3275-3286.

Kwon, M.J., Boyanov, M.I., Antonopoulos, D.A., Brulc, J.M., Johnston, E.R., Skinner, K.A., Kemner, K.M., and O’Loughlin, E.J. (2014). Effects of dissimilatory sulfate reduction on Fe^III^ (hydr)oxide reduction and microbial community development. Geochimica et Cosmochimica Acta 129, 177-190.

Kwon, M.J., O’Loughlin, E.J., Boyanov, M.I., Brulc, J.M., Johnston, E.R., Kemner, K.M., and Antonopoulos, D.A. (2016). Impact of organic carbon electron donors on microbial community development under iron- and sulfate-reducing conditions. PLOS ONE 11, e0146689.

Lauber, C.L., Hamady, M., Knight, R., and Fierer, N. (2009). Pyrosequencing-based assessment of soil pH as a predictor of soil bacterial community structure at the continental scale. Applied and Environmental Microbiology 75, 5111-5120.

Laverman, A.M., Garnier, J.A., Mounier, E.M., and Roose-Amsaleg, C.L. (2010). Nitrous oxide production kinetics during nitrate reduction in river sediments. Water Research 44, 1753-1764.

Lebuhn, M., Achouak, W., Schloter, M., Berge, O., Meier, H., Barakat, M., Hartmann, A., and Heulin, T. (2000). Taxonomic characterization of *Ochrobactrum* sp. isolates from soil samples and wheat roots, and description of *Ochrobactrum tritici* sp. nov. and *Ochrobactrum grignonense* sp. nov. International Journal of Systematic and Evolutionary Microbiology 50, 2207-2223.

Leonard, S., Hommais, F., Nasser, W., and Reverchon, S. (2017). Plant–phytopathogen interactions: bacterial responses to environmental and plant stimuli. Environmental Microbiology 19, 1689-1716.

Lessie, T.G., and Phibbs, P.V. (1984). Alternative pathways of carbohydrate utilization in pseudomonads. Annual Review of Microbiology 38, 359-388.

Luo, F., Gitiafroz, R., Devine, C.E., Gong, Y., Hug, L.A., Raskin, L., and Edwards, E.A. (2014). Metatranscriptome of an anaerobic benzene-degrading, nitrate-reducing enrichment culture reveals involvement of carboxylation in benzene ring activation. Applied and Environmental Microbiology 80, 4095-4107.

Ma, Y., Wang, L., and Shao, Z. (2006). *Pseudomonas*, the dominant polycyclic aromatic hydrocarbon-degrading bacteria isolated from Antarctic soils and the role of large plasmids in horizontal gene transfer. Environmental Microbiology 8, 455-465.

McMurdie, P.J., and Holmes, S. (2013). phyloseq: An R package for reproducible interactive analysis and graphics of microbiome census data. PLOS ONE 8, e61217.

Meyer, F., Paarmann, D., D’Souza, M., Olson, R., Glass, E.M., Kubal, M., Paczian, T., Rodriguez, A., Stevens, R., Wilke, A., Wilkening, J., and Edwards, R.A. (2008). The metagenomics RAST server - a public resource for the automatic phylogenetic and functional analysis of metagenomes. BMC Bioinformatics 9, 386.

Nemergut, D.R., Schmidt, S.K., Fukami, T., O’Neill, S.P., Bilinski, T.M., Stanish, L.F., Knelman, J.E., Darcy, J.L., Lynch, R.C., Wickey, P., and Ferrenberg, S. (2013). Patterns and processes of microbial community assembly. Microbiology and Molecular Biology Reviews 77, 342-356.

O’Brien, S.L., Gibbons, S.M., Owens, S.M., Hampton-Marcell, J., Johnston, E.R., Jastrow, J.D., Gilbert, J.A., Meyer, F., and Antonopoulos, D.A. (2016). Spatial scale drives patterns in soil bacterial diversity. Environmental Microbiology 18, 2039-2051.

Onesios-Barry, K.M., Berry, D., Proescher, J.B., Sivakumar, I.K.A., and Bouwer, E.J. (2014). Removal of pharmaceuticals and personal care products during water recycling: microbial community structure and effects of substrate concentration. Applied and Environmental Microbiology 80, 2440-2450.

Ottesen, E.A., Young, C.R., Gifford, S.M., Eppley, J.M., Marin, R., Schuster, S.C., Scholin, C.A., and DeLong, E.F. (2014). Multispecies diel transcriptional oscillations in open ocean heterotrophic bacterial assemblages. Science 345, 207-212.

Peralta, A. L., Matthews, J.W., and Kent, A.D. (2010). Microbial community structure and denitrification in a wetland mitigation bank. Applied and Environmental Microbiology 76, 4207-4215.

Prosser, J.I. (2015). Dispersing misconceptions and identifying opportunities for the use of ‘omics’ in soil microbial ecology. Nature Reviews Microbiology 13, 439-446.

Raskin, L., Rittmann, B.E., and Stahl, D.A. (1996). Competition and coexistence of sulfate-reducing and methanogenic populations in anaerobic biofilms. Applied and Environmental Microbiology 62, 3847-3857.

Robinson, M.D., McCarthy, D.J., and Smyth, G.K. (2010). edgeR: a Bioconductor package for differential expression analysis of digital gene expression data. Bioinformatics 26, 139-140.

Roesch, L.F.W., Fulthorpe, R.R., Riva, A., Casella, G., Hadwin, A.K.M., Kent, A.D., Daroub, S.H., Camargo, F.A.O., Farmerie, W.G., and Triplett, E.W. (2007). Pyrosequencing enumerates and contrasts soil microbial diversity. The ISME Journal 1, 283-290.

Salles, J.F., van Veen, J.A., and van Elsas, J.D. (2004). Multivariate analyses of *Burkholderia* species in soil: effect of crop and land use history. Applied and Environmental Microbiology 70, 4012-4020.

Shade, A., Hogan, C. S., Klimowicz, A.K., Linske, M., McManus, P. S., and Handelsman, J. (2012). Culturing captures members of the soil rare biosphere. Environmental Microbiology 14, 2247-2252.

Spiers, A.J., Buckling, A., and Rainey, P.B. (2000). The causes of *Pseudomonas* diversity. Microbiology 146, 2345-2350.

Stanier, R.Y., Palleroni, N.J., and Doudoroff, M. (1966). The aerobic pseudomonads: a taxonomic study. Microbiology 43, 159-271.

Stegen, J.C., Lin, X., Fredrickson, J.K., Chen, X., Kennedy, D.W., Murray, C.J., Rockhold, M.L., and Konopka, A. (2013). Quantifying community assembly processes and identifying features that impose them. The ISME Journal 7, 2069-2079.

Stegen, J.C., Lin, X., Konopka, A.E., and Fredrickson, J.K. (2012). Stochastic and deterministic assembly processes in subsurface microbial communities. The ISME Journal 6, 1653-1664.

Stopnisek, N., Bodenhausen, N., Frey, B., Fierer, N., Eberl, L., and Weisskopf, L. (2014). Genus-wide acid tolerance accounts for the biogeographical distribution of soil *Burkholderia* populations. Environmental Microbiology 16, 1503-1512.

Sutton, N.B., van Gaans, P., Langenhoff, A.A.M., Maphosa, F., Smidt, H., Grotenhuis, T., and Rijnaarts, H.H.M. (2013). Biodegradation of aged diesel in diverse soil matrixes: Impact of environmental conditions and bioavailability on microbial remediation capacity. Biodegradation 24, 487-498.

Tiedje, J.M., Asuming-Brempong, S., Nüsslein, K., Marsh, T.L., and Flynn, S.J. (1999). Opening the black box of soil microbial diversity. Applied Soil Ecology 13, 109-122.

Ushiba, Y., Takahara, Y., and Ohta, H. (2003). *Sphingobium amiense* sp. nov., a novel nonylphenol-degrading bacterium isolated from a river sediment. International Journal of Systematic and Evolutionary Microbiology 53, 2045-2048.

Weber, K.P., and Legge, R.L. (2009). One-dimensional metric for tracking bacterial community divergence using sole carbon source utilization patterns. Journal of Microbiological Methods 79, 55-61.

Whitman, W.B., Coleman, D.C., and Wiebe, W.J. (1998). Prokaryotes: The unseen majority. Proceedings of the National Academy of Sciences 95, 6578-6583.

Yabuuchi, E., Kosako, Y., Oyaizu, H., Yano, I., Hotta, H., Hashimoto, Y., Ezaki, T., and Arakawa, M. (1992). Proposal of *Burkholderia* gen. nov. and transfer of seven species of the genus *Pseudomonas* Homology Group II to the new genus, with the type species *Burkholderia cepacia* (Palleroni and Holmes 1981) comb. nov. Microbiology and Immunology 36, 1251-1275.

Yabuuchi, E., Kosako, Y., Yano, I., Hotta, H., and Nishiuchi, Y. (1995). Transfer of two *Burkholderia* and an *Alcaligenes* species to *Ralstonia* gen. nov. Microbiology and Immunology 39, 897-904.

Youngblut, N.D., Dell’Aringa, M., and Whitaker, R.J. (2014). Differentiation between sediment and hypolimnion methanogen communities in humic lakes. Environmental Microbiology 16, 1411-1423.

Zeglin, L.H. (2015). Stream microbial diversity in response to environmental changes: review and synthesis of existing research. Frontiers in Microbiology 6, 1-15.

Zhang, Y., Zheng, Y., Hu, J., Du, N., and Chen, F. (2014). Functional diversity of the microbial community in healthy subjects and periodontitis patients based on sole carbon source utilization. PLOS ONE 9, e91977.

